# Comparative analysis of gene regulatory networks identifies conserved regulators in seed plants

**DOI:** 10.1101/2023.11.20.567877

**Authors:** Donat Wulf, Andrea Bräutigam

## Abstract

Gene regulatory networks based on transcription factors control development and environmental responses in plants. Networks calculated by the machine learning algorithm random forest decision tree-based regression for the grasses barley, maize, wheat, Brachypodium, sorghum, and rice compared with Arabidopsis and an alga show substantial conservation. The degree of conservation depends on phylogenetic closeness. The processes, which are conserved between all species include basic cellular functions while the processes conserved in the grasses also more specific gene ontology terms. In the three species with a carbon concentration mechanism, photorespiration is partially disassociated from photosynthetic regulation. In contrast, in the C4 species, the regulation of C4 genes associates with photosynthetic regulation. The comparative analyses reveal conserved transcription factors, which control photosynthesis in seed plants but not in the alga. An analysis pipeline for the general transfer of information between the small weed Arabidopsis and the commercially relevant grasses is presented.

## Introduction

Progress in gene function identification in plant research has mostly occurred with the small weed *Arabidopsis thaliana* which has no importance as a crop plant. In contrast, wheat provides 20% of calories produced. Together with other grasses such as rice (12%), barley (5%), sorghum (2%) and maize (11%), grasses account for 50% of calories produced (D’Odorico et al., 2014). Information transfer between Arabidopsis and grasses was successful in the past (Pucker et al., 2020; Shi et al., 2020; Hughes and Langdale, 2022; Busche et al., 2023) and is highly desirable for conceptualizing breeding and gene editing of plants. Arabidopsis and the grasses share a common ancestor 150 million years ago (Chaw et al., 2004). The gene regulatory networks underlying developmental programs and responses to biotic and abiotic challenges are likely at least partially conserved.

In the case of gene regulation, information transfer between species is particularly difficult. Plant species have undergone partial and whole genome duplications (WGDs) along different branches of the phylogenetic tree which leads to redundancy (Wagner, 1996; Martín et al., 2016; Wang et al., 2013). WGDs in polyploids influence motif frequencies and network motifs (Almeida-Silva and Van de Peer, 2023). Between the last common ancestor of Arabidopsis and grasses three WGD events occurred towards Arabidopsis, four WGDs occurred towards *Zea mays*, and three towards rice and barley (Van de Peer et al., 2017). Several important crop genomes show recent polyploidy such as wheat with three constituent genomes (IWGSC et al., 2018) and oilseed rape with two constituent genomes (Lee et al., 2020). Plant breeding also had an influence on the genomes of crop plants (Jayakodi et al., 2020). WGDs are followed by at least partial diploidization in which some but not all genes are lost again (Kuzmin et al., 2022). Transcription factors are preferably retained upon genome duplication (Wendel, 2000) and over time this leads to a divergence in expression pattern (Blanc and Wolfe, 2004). This makes the identification of functional orthologues for transcription factors based on sequence more difficult than identifying orthologues of genes in primary metabolism. To overcome this obstacle, it is necessary to compare expression patterns across different species. Such transcriptomic studies which are comparable between different species exist, but only cover a limited number of conditions (Bräutigam et al., 2011; Vercruysse et al., 2020; Julca et al., 2021). Furthermore, plants live in a wide range of different ecological niches, which makes finding comparable sampling conditions difficult (Wilkins et al., 2016; Chaloner et al., 2020).

Network inference with the random forest decision tree-based algorithm GENIE3 (Huynh-Thu et al., 2010) was used to infer a gene regulatory network (GRN) in wheat (Ramírez-González et al., 2018) and analyze polyploid evolution (Almeida-Silva and Van de Peer, 2023). Recently, GENIE3 was used for identification and validation of regulators involved in photosynthesis in *A. thaliana* (Halpape et al., 2023) and in the regulation of photoprotection and the carbon concentrating mechanism in *Chlamydomonas reinhardtii* (Arend et al., 2023; Wulf et al., 2023) making it a promising candidate to compare gene regulatory networks across species. By performing GRN inference based on expression data comparability of individual samples is not necessary, because transcription factors are examined in relation to all other genes. Machine learning based analysis is dependent of a high amount input data. Therefore, we focused on economically important crop plants (D’Odorico et al., 2014) *Brachypodium distachyon*, *Hordeum vulgare*, *Oryza sativa*, *Sorghum bicolor*, *Triticum aestivum* and *Z. mays* with sufficient data in the sequence read archive. For these species a wide range of transcriptome data from different conditions is available (Katz et al., 2022). The grass species represent the *Andropogonae* in the PACMAD clade with *Z. mays* and *S. bicolor*, the Pooidae with *B. distachyon*, *H. vulgare* and *T. aestivum* in the BEP clade, and the Ehrhartoidae with *O. sativa* also in the BEP clade (Christin et al., 2009). The split between BEP and PACMAD can be roughly dated at 55 million years ago (Christin et al., 2014). By including previously published and validated gene regulatory networks of *A. thaliana* (Halpape et al., 2023) and *C. reinhardii* (Wulf et al., 2023) the analysis is extended to dicots and algae to include more distant species at 150 million years between eudicots and monocots and 950 million years between the chlorophyta and the streptophyta (Magallón et al., 2013; Hedges et al., 2004). We hypothesized that the comparison of networks will identify conserved and divergent regulatory events.

The photosynthetic trait was chosen for comparison. It is always indispensable for photoautotrophic growth and therefore likely under selection. The presence of carbon concentration mechanisms provides the opportunity to test if metabolic pathways associated with photosynthesis are associated with its regulon. We hypothesized that transcriptional regulation of photosynthesis is largely conserved. The transcription factors GOLDEN2-LIKE (GLK) were originally identified in *Z. mays* and shown to control photosynthetic transcripts in the monocot *Z. mays*, the dicot *A. thaliana*, and the liverwort *Marchantia polymorpha* (Fitter et al., 2002; Yasumura et al., 2005; Yelina et al., 2023) already demonstrating a large degree of conservation. The species under investigation operate different modes of photosynthesis. *B. distachyon*, *H. vulgare*, *O. sativa*, and *T. aestivum* operate C3 photosynthesis. One metabolic shortcoming of C3 photosynthesis is RuBisCO. It not only fixes carbon but also oxygen. The oxygenation reaction results in phosphoglycolate, which has to be detoxified by photorespiration (PR), a metabolic pathway associated with photosynthesis. This pathway consumes 2 NADPH and 3.5 ATP (Walker et al., 2016). Furthermore, CO_2_ is released making this costly for the plant. *C. reinhardtii*, *S. bicolor* and *Z. mays* operate a photosynthesis associated pathway, the carbon concentrating mechanism, which eliminates the need for high flux photorespiration (Wang et al., 2015; Mallmann et al., 2014). The photorespiratory pathway is associated with the photosynthetic regulon in C3 plants such as Arabidopsis (Halpape et al., 2023). In *C. reinhardtii*, CO_2_ is converted to H_3_O^-^ and transported into the chloroplast into the pyrenoid, which is structurally made out of a starch shell and condensed RuBisCO (Wang et al., 2015). *S. bicolor* and *Z. mays* both operate the C4 cycle (Sage, 2004). Here, CO_2_ is shuttled via a C4 acid into the bundle sheath where it is then fixated by RuBisCO. Genes belonging to the C4 cycle are genes of the primary metabolism of plants and are therefore present in C3 plants (Aubry et al., 2011). Association of C4 gene regulation to the photosynthetic regulon has been suggested (Külahoglu et al., 2014; Aubry et al., 2016). We hypothesized that in C4 species, the C4 genes are associated with the photosynthetic regulon while photorespiratory genes likely diminish in association.

We show that GRNs of similar information content can be constructed for all species under investigation. The GRNs are conserved between species to varying degrees and the degree of conservation depends on phylogenetic distance. Photosynthesis is indeed a conserved process with orthologous transcription factors (TFs) identified as its regulators. We show that evolutionary changes in photosynthetic mode have concomitant changes in the regulatory network. Finally, the comparative method can be extended to agronomically relevant traits of choice, i.e. analyses of biotic interactions.

## Material and Methods

### Genome versions

### Network inference

The sequence read archive (Katz et al., 2022) was queried for RNA-seq experiments for *B. distachyon*, *H. vulgare*, *O. sativa*, *S. bicolor* (“Species name” AND “RNA”). By manual curation wild type studies were selected (Table S1). The raw read files were downloaded with fasterq-dump (v2.11.2) and mapped on the primary transcriptome of the respective species with Kallisto (v0.46.1) (Bray et al., 2016) with an average library length of 200 and a standard deviation of 20 for single-end reads and default parameters for paired-end reads. Experiments were filtered for a mapping rate of at least 70% and at least 7.5 million mapped reads to ensure sufficient depth for analysis of regulatory events. The resulting expression matrix is publicly available as a data publication (https://gitlab.ub.uni-bielefeld.de/computationalbiology/comp_net; DOI to be obtained upon acceptance to ensure revisions are reflected in the git).

Arabidopsis transcription factors were obtained from Halpape et al. (2023) and *Z. mays* transcription factors were obtained from TAP-scan (Petroll et al., 2021) (Table S2). Transcription factors were transferred to the other species based on Plaza 4.5 (Van Bel et al., 2018). Gene regulatory networks were inferred with the random forest decision tree regression based algorithm GENIE3 (Huynh-Thu et al., 2010) with 1,000 decision trees and the square root of regulators. Gene Ontology (GO) terms were transferred based on the best blastp (Altschul et al., 1990) hit from *A. thaliana* to the other species with a blastp (e < 1e-5) to exclude annotation artefacts introduced by different GO annotations. Genes with a weight greater than 0.005 were considered as target of the transcription factor (Arend et al., 2023; Halpape et al., 2023; Wulf et al., 2023). For each target list of the transcription factors a GO term enrichment was performed with topGO (v2.52.0) (Alexa and Rahnenführer, 2023) using the classic fisher method for the ontology biological process with a significance cut-off at p-value < 1e-10 (Table S5).

### Comparison of orthologue regulators

To assign orthologue groups orthofinder2 (Emms and Kelly, 2015) was used on the primary proteomes of *A. thaliana*, *B. distachyon*, *C. reinhardtii*, *O. sativa*, *H. vulgare*, *S. bicolor*, *T. aestivum*, and *Z. mays*. To test whether two TFs share a significant amount of target orthogroups a fishers exact test was performed. An orthologue gene group was considered regulated by both regulators if one gene of each species was a target gene of the respective regulator. Orthologous regulators whose targets overlapped significantly (p-value < 1e-10) were considered conserved. Orthologue regulator groups were considered conserved if one pairwise comparison was found to be conserved. For visualization the minimum –log_10_(p-value) was plotted capped at 1e-100. The GO term was considered conserved between two orthologue transcription factors if both transcription factors had a significant GO term enrichment (p-value < 1e-10) in the same GO term and a significant overlap in the target genes. This method tests if the regulon overall is conserved.

### Identification of light harvesting complexes and photosystems, Calvin–Benson–Bassham cycle, photorespiration and C4 cycle genes

Light harvesting complexes and photosystems, Calvin–Benson–Bassham cycle, and photorespiration genes were obtained from Halpape et al. (2023) and transferred by sequence homology with a blastp (Altschul et al., 1990) search with a p-value cut-off of 1e-5 to the other species used in this study (Table 1). The C4 cycle genes were obtained from Schlüter and Weber (2016) and searched against *Z. mays* with a blastp (Altschul et al., 1990) search with a p-value cut-off of 1e-5. The highest expressed leaf isoform was selected based on Sekhon et al. (2011). These genes were then searched in the other species (Table 1) with blastp (Altschul et al., 1990) with a p-value cut-off of 1e-5. For *T. aestivum*, the best blast hit from the A, C and D genome was used. For all species the photosynthetic genes are available in Table S3.

**Table 1:** Species and genome versions used in this study.

| Species | Genome version |
| --- | --- |
| *Arabidopsis thaliana* | TAIR10 |
| *Chlamydomonas reinhardtii* | v5.6 |
| *Brachypodium distachyon* | v3.1 |
| *Oryza sativa* | v7.0 |
| *Hordeum vulgare* | r1 |
| *Sorghum bicolor* | v3.1 |
| *Triticum aestivum* | iwgsc v1.0 |
| *Zea mays* | APGv4 |

### Identification of photosynthesis and C4 photosynthesis related regulators

For each transcription factor of each network a fishers exact test among target genes of each TF for light harvesting complexes and photosystems (LHC & PS), Calvin–Benson–Bassham cycle (CBBc), photorespiration (PR) and C4 cycle was performed. Regulators were assigned to pathways based on a p-value smaller than 1e-10 (Table S5). Eigenvalue centrality was calculated with the bonacich power centrality scores (Bonacich, 1987) between the regulators and the LHC & PS, CBBc, PR, and C4 genes using all weights from the sub networks. Conserved photosynthesis regulators were identified by comparing their best blastp hit to *A. thaliana* with a p-value cut-off of 1e-5.

### Visualization and used R libraries

Data analysis was performed in R v4.3.1 using tidyverse, sna, and topGO. Visualisations were done with ggplot2, ggupset, cowplot, and igraph,

## Results

The number of available wild type experiments for the eight species under investigation ranged between 769 in *C. reinhardtii* and 6033 in *A. thaliana* because of different numbers of available experiments on NCBI. The number of transcription factors ranged between 237 in *C. reinhardtii* and 7400 in *T. aestivum* and reflects the number of WGDs, small scale duplications, and the degree of diploidization which occured. *Z. mays* had 3200 annotated transcription factors and the other species contained around 2300 annotated transcription factors (Table 2). The expression matrix for each species enabled the comparison of overview metrics. For most species the number of genes with a tpm > 1 in at least one condition ranged from 95.6% in *B. distachyon* to 99.8% in *A. thaliana*. The outlier was T. aestivum with 70.8% of genes with a tpm of at least one (Table 2). For the transcription factor this was slightly higher for all species. *T. aestivum saw* the strongest increase to 96.9% (Table 2). By defining housekeeping genes with a coefficient of variation below 50 we were able to identify housekeeping genes. The amount ranged between 13% for *B. distachyon* and 2% for *C. reinhardii* and *Z. mays* (Table 2). 5% of all transcription factors were housekeeping genes compared to 3% of non-transcription factors. 0.5% of photosynthesis genes were among the house keeping genes.

**Table 2:** Number of regulators, number of mapped wild type experiment genes with a coefficient of variation smaller than 50 and the frequency of genes with a coefficient of variation smaller than 50 for the species A. thaliana, B. distachyon, C. reinhardtii, H. vlgare, O. sativa, S. bicolor, T. aestivum, and Z. mays.

| Species | Number of regulators | Number of wild type experiments | Percentage expressed genes | Percentage expressed TFs | Genes with cov<50 | Frequency of genes with cov<50 |
| --- | --- | --- | --- | --- | --- | --- |
| *A. thaliana* | 2402 | 6033 | 99.8% | 100% | 3747 | 0.07 |
| *B. distachyon* | 2299 | 802 | 95.6% | 97.4% | 4108 | 0.08 |
| *C. reinhardtii* | 237 | 789 | 99.0% | 100% | 273 | 0.01 |
| *H. vulgare* | 2170 | 1089 | 95.0% | 97.3% | 1175 | 0.01 |
| *O. sativa* | 2315 | 1415 | 97.5% | 99.5% | 1445 | 0.01 |
| *S. bicolor* | 2366 | 1090 | 96.6% | 98.6% | 4572 | 0.10 |
| *T. aestivum* | 7400 | 769 | 70.8% | 96.9% | 8748 | 0.02 |
| *Z. mays* | 3248 | 2022 | 96.9% | 99.0% | 759 | 0.01 |

### Compare covariation of TFs to other genes

For every TF in each inferred GRN a GO term enrichment was calculated. The information content represented by the number of transcription factors with a significant (p < 1^-10^) enrichment varied between the different networks. In *T. aestivum* in absolute terms the most TFs (3313) have a significant GO term enrichment. The smallest number of TFs with an enrichment was identified in *C. reinhardtii* with 237. The number of TFs with enrichment was between 853 in *B. distachyon* and 1083 in *Z. mays*. In *A. thaliana*, 1274 TFs had a significant enrichment (Figure 1A). To compare the quality of the GRNs, we used three measures. In addition to the frequency of TFs with a GO term enrichment, we used the number of unique enriched GO terms overall and the number of unique enriched GO terms with the lowest p-values of the enrichment for a TF (Figure 1B, C, D). The frequency of TFs with an enrichment varied between 0.53 in *A. thaliana* and 0.33 in *Z. mays*. In *B. distachyon*, *S. bicolor*, *O. sativa*, *and T. aestivum*, the frequency was between 0.37 and 0.45.

**Figure 1:**
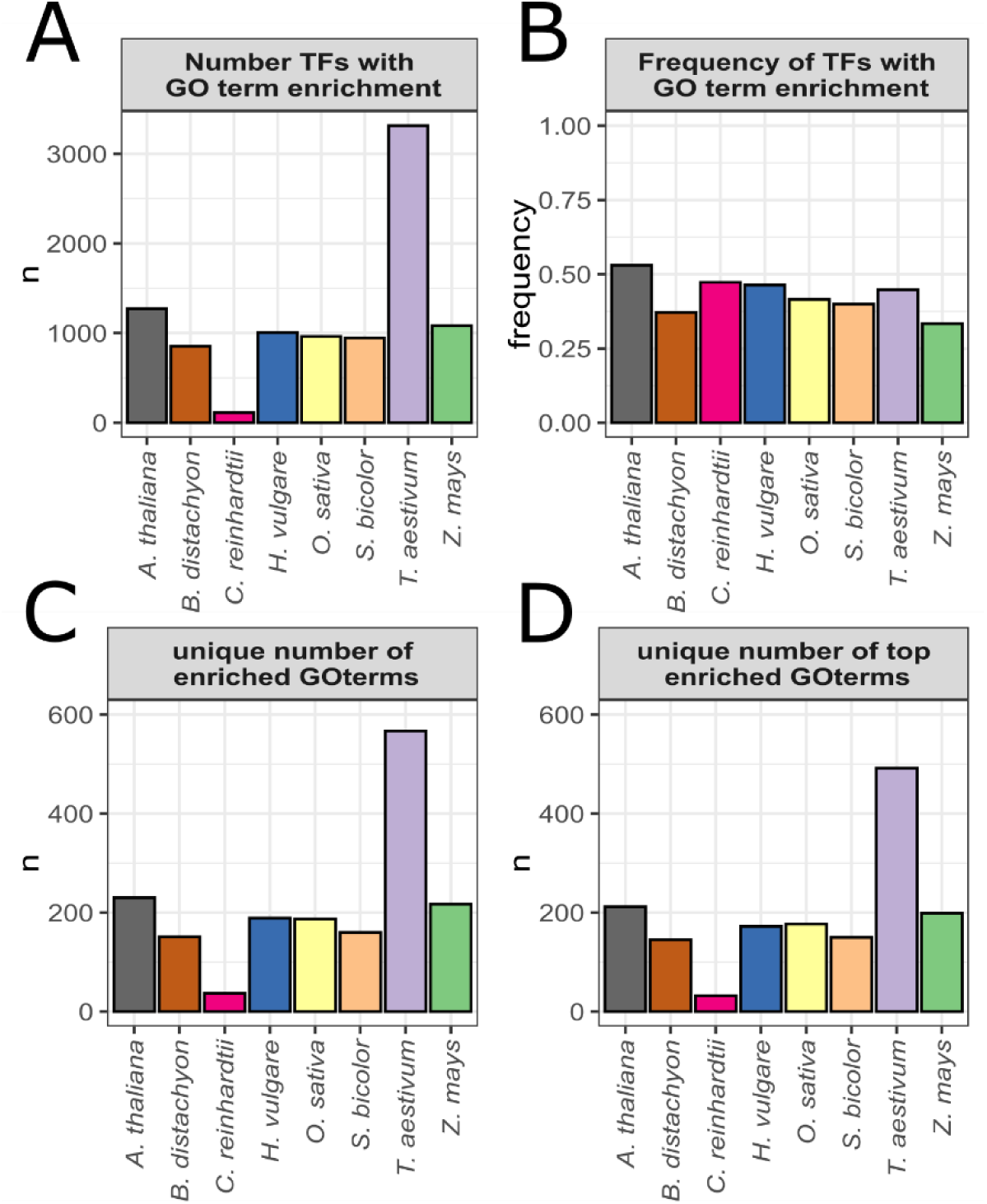
GO term stats of inferred GRNs. A: Number of transcriptions factors with a GO term enrichment p < 1^-10^. B: Frequency of transcription factors with a GO term enrichment p < 1^-10^. C: Unique number of enriched GO terms p < 1^-10^. D: Unique number of enriched GO terms with the highest enrichment p < 1^-10^. A. thaliana (black), B. distachyon (brown), C. reinhardtii (pink), H. vulgare (blue), O. sativa (yellow), S. bicolor (orange), T. aestivum (purple), and Z. mays (green).

*H. vulgare* and *C. reinhardtii* reached a frequency of 0.46 and 0.47 (Figure 1C). To estimate how many processes the GRNs represent, we examined the number of different GO terms with a significant enrichment (p < 1^-10^) in each network. In *C. reinhardtii*, 37 different GO terms had a significant enrichment. In *T. aestivum*, 567 different GO terms had a significant enrichment. In *A. thaliana* and the remaining species, 230 to 160 different GO terms had a significant enrichment (Figure 1C). How good these processes are separated across different TFs was tested by only counting the strongest significantly (p < 1^-10^) enriched GO terms for each transcription factor. A slight reduction of at most 14% in unique GO terms was observed compared to all significant GO terms (Figure 1C, D). Overall, the information content of the networks was reasonably similar with 33% to 50% TFs having enrichments when one considers that the amount of data and the type of data underlying each network is widely divergent (Table S5) For comparison, pairwise conserved orthologous groups of the regulators of the different GRNs were identified. The number of conserved orthogroups on the sequence level serves as the baseline.

The number of orthogoups conserved on sequence level ranged between 1500 and 1177 for the grasses. Between *A. thaliana* and the grasses, the number of orthogroups ranged between 908 (*S. bicolor*) and 764 (*H. vulgare*) and between *C. reinhardtii* and the monocots and *A. thaliana* ranged between 64 and 59. An orthogroup was considered conserved in the gene regulatory network if one pairwise comparison yielded a p-value smaller than 1^-10^. For plotting, *A. thaliana* as a representative of the eudicots was used for reference, *Z. mays* was used as a representative of the monocots, and *C. reinhardtii* was used as a representative of the alga.

Boxplots show the degree of conservation between all species and *A. thaliana* (Figure 2A) representing 150 million years since divergence and 950 million years since divergence. A small subset of TFs shows extremely high conservation as they are outliers to the top of the boxplot. All species except *C. reinhardtii* reach a conservation frequency of 38% or above. In the alga, 21% of TFs have detectable conserved function compared to Arabidopsis (Figure 2A middle panel). Relative to *A. thaliana*, the grass species show a frequency of conservation with values between 46% for *O. sativa* and 38% for *T. aestivum*. As a non-threshold measure for conservation we also analyzed the p-value of the median transcription factor. The median of the p-value resulting from the fisher test of the overlaps ranged between 1.8e^-6^ for *T. aestivum* and 8.7e^-10^ for *O. sativa* compared to 3.9e^-3^ for *C. reinhardtii*. The quantiles follow the same pattern (Figure 2A top panel). The observed degree of conservation is a lower bound given that the networks are calculated with varying amounts of data and varying experiments represented. The plot based on *Z. mays* offers a grass-centric view (Figure 2B) and represents 55, 150 and 950 million years since divergence. The frequency of conserved regulator orthogroups was highest with 64% to *S. bicolor* followed by *O. sativa, H. vulgare, B. distachyon,* and *T. aestivum* with 48%. Among the grasses conservation is higher than in *A. thaliana* with 42% and than in *C. reinhardtii* with only 14%. The order of frequency of conserved orthogroups follows phylogeny. The highest conservation is detected between *Z. mays* and *S. bicolor* which are both Andropogonae in the PACMAD branch of grasses (Christin et al., 2009) (Figure 2B middle panel). The degree of conservation is also higher between *Z. mays* and the other grass species with a median of the p-value resulting from the fisher test of the overlaps of 4.3e^-10^ in *T. aestivum* and 6.6e^-10^ *S. bicolor* (Figure 2A, B top panel). The plot based on *C. reinhardtii* offers the most distant comparison representing 950 million years of evolution in all cases (Figure 2C). The frequency of conserved overlaps is between 11% in *B. distachyon* and 21% in *A. thaliana* and therefore much lower as compared to the frequency of conservation of *A. thaliana* and *Z. mays* to the other plants (Figure 2A, B, C, middle panel). The median p-value of the overlaps is between 0.02 in *O. sativa* and 0.004 in *A. thaliana*. The best p-value for the *C. reinhardtii* overlap was to *A. thaliana* with 5.6^-67^ (Figure 2C top panel).

**Figure 2:**
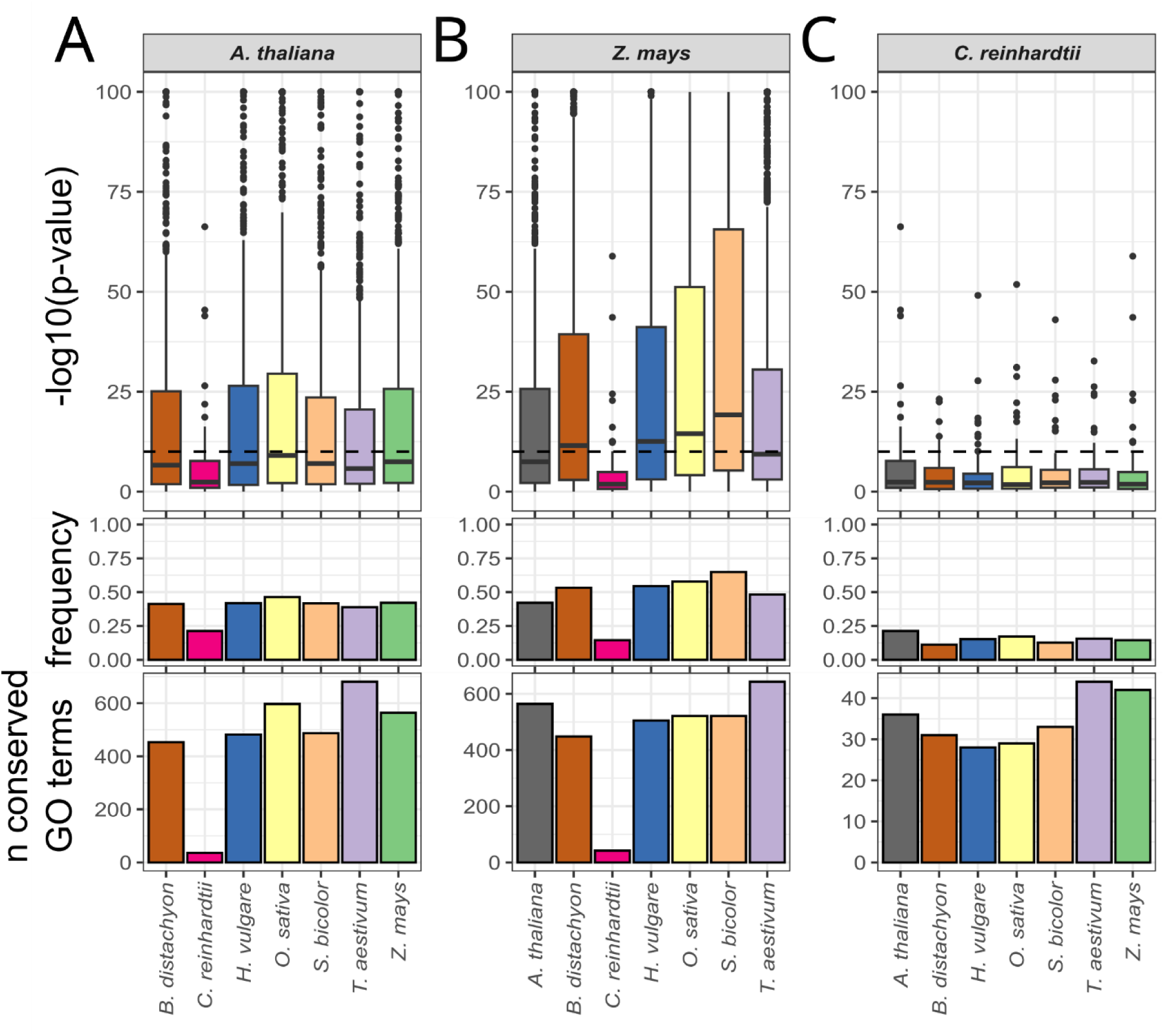
Conservation of orthologue regulators (top), frequency of conserved orthogroups (middle) and number of conserved GO terms (bottom) compared to A. thaliana (A), Z. mays (B) and C. reinhardtii of A. thaliana (black), B. distachyon (brown), C. reinhardtii (pink), H. vulgare (blue), O. sativa (yellow), S. bicolor (orange) T. aestivum (purple) and Z. mays (green). Y-axis are scaled differently in the bottom pannel for better visibilty.

The number of conserved GO terms between species is another measure of similarity. The types of functions represented by those GO terms show, which processes are conserved during evolution. To estimate which and how many processes are regulatory conserved between the different species, we examined the overlap between significantly enriched GO terms of conserved orthologue regulator groups. Over all networks, 2384 GO terms had a significant enrichment. Viewed from the dicot *A. thaliana*, 37 processes were regulatory conserved to *C. reinhardtii* (Figure 2A bottom panel). Toward the grasses, this ranged from 453 in *B. distachyon* to 681 in *T. aestivum*. The 37 processes conserved toward the alga were regulatory conserved over all species. These processes were associated with general cell function e.g., translation, peptide biosynthetic process, nucleic acid metabolic process, and response to light stimulus (Table S4). Toward all monocots, 353 different processes were conserved overall. Within these processes are also more specific GO terms e.g. cell wall polysaccharide biosynthetic process, carotenoid metabolic process, pollen tube development, response to heat, and photosystem II assembly (Table S4). Viewed from the Andropogonae monocot *Z. mays*, the number of conserved processes in the grasses ranged from 448 in *B. distachyon* to 643 in *T. aestivum*. Toward *A. thaliana*, 564 more processes were conserved to *Z. mays* than in all grasses except *T. aestivum*, but the overlap between *Z. mays* and the other grasses is larger than in *A. thaliana* (Figure 2A, B bottom panel). Comparing *Z. mays* to the monocots results in 358 conserved regulatory processes. These were 34 more than in the comparison if *A. thaliana* was included. Within these monocot specific conserved regulatory processes are e.g. shoot development, defense response to bacterium, secondary metabolite biosynthetic process, response to hormone, and response to *oomycetes* (Table S4) and therefore both response terms and constitutive terms are included. Viewed from *C. reinhardtii*, the least amount of conserved regulatory processes is found. It ranged between 28 conserved processes in *H. vulgare* and 44 to *T. aestivum* (Figure 2C). If fuzzily conserved processes are allowed in the comparison to *C. reinhardtii*, we identify cell cycle, DNA repair, and DNA replication as conservedly regulated in 4 out of 7 other species (Table S6).

Photosynthesis represents a pathway with high likelihood for conservation. Regulators for photosystems and light harvesting complexes (PS&LHCs), Calvin Benson Bassham cycle (CBBc), photorespiration (PR), and C4 photosynthesis (C4) were identified by enrichment analysis for genes involved in these pathways (Table S5). The number of identified regulators differed for each pathway and each species. The numbers ranged between 18 for *C. reinhardtii* and 96 for *T. aestivum*. For *A. thaliana*, *B. distachyon, H. vulgare*, and *O. sativa*, regulators above the significance threshold of p < 1e-10 were identified for PS&LHCs, CBBc and PR, but not for C4 genes (Figure 3, Figure S1). In *C. reinhardtii*, only regulators for the photosystems and LHCs were identified (Figure 3B) but no regulators for the other components crossed the threshold of p < 1e-10. In *Z. mays* and *S. bicolor*, no regulators above the threshold were identified for PR but regulators were identified for C4 cycle genes (Figure 3D, Figure S1). In *T. aestivum*, regulators for all processes were identified, but the number of regulators for C4 cycle genes was five (Figure S1, Table S5). For *A. thaliana*, *B. distachyon*, *H. vulgare, O. sativa*, and *T. aestivum*, the PR regulators were for the most part shared with CBBc regulators (Figure S1, Table S5). These data are dependent on a threshold cut-off. To test how strong the identified regulators are connected to the different domains of photosynthesis, we used a cut-off independent centrality measure. For calculation, the eigenvalue between each transcription factor with a photosynthesis enrichment and the LHC&PS, CBBc, PR, and C4 genes was calculated. For the Photosystems and LHC, the centrality was high for all species. For the CBBc, a reduction in the eigenvalue centrality was observed in *C. reinhardtii*. The photorespiratory genes were strongly connected to the photosynthesis regulators only in C3 species, but a decrease in the median was observed for *C. reinhardtii*, *S. bicolor*, and *Z. mays*. All these species contain a CCM which reduces the flux through photorespiration. Regulation of photorespiration at least partially disconnects from regulation of photosynthesis in species with a CCM (Figure 3E). The C4 cycle genes were only central in the network in the C4 species *S. bicolor* and *Z. mays*. In the C3 species including *C. reinhardtii*, the centrality of these genes was lower than in the C4 species (Figure 3E). Hence, the C4 cycle is only connected to the photosynthesis regulon in species in which genes carry a flux vital for efficient photosynthesis. The evolution of photosynthesis associated pathways and their phenotypes can thus be traced in the GRNs.

**Figure 3:**
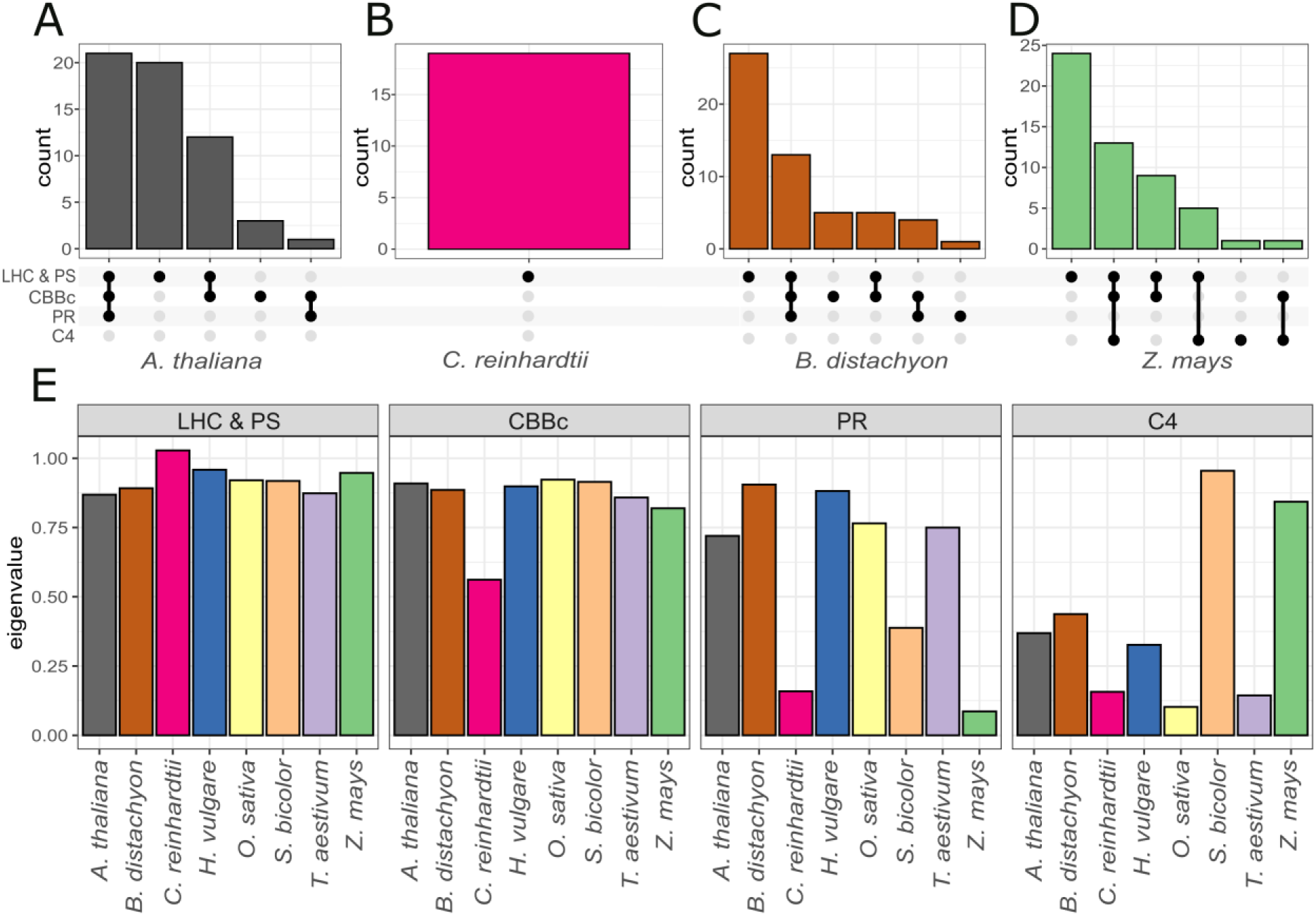
Shared regulators between photosynthetic processes of A. thaliana, C. reinhardtii, B. distachyon and Z. mays in the processes photosystems (PS), Calvin Benson Bassham cycle (CBBc), photorespiration (PR), and C4 cycle (C4). Eigenvalue centrality measure of PS, CBBc, PR and C4 cycle genes in A. thaliana, B. distachyon, C. reinhardtii, H. vulgare, O. sativa, S. bicolor, T. aestivum, and Z. mays.

The substantial degree of conservation among the seed plants should enable detection of conserved regulators. To identify these conserved photosynthesis regulators, the predicted photosynthesis regulators (Figure 4, Figure S1, Table S7) were tested for sequence similarity by grouping them based on their best blast hit with *A. thaliana*. *C. reinhardtii* had a conserved enrichment in photosynthesis related genes with *Z. mays* for ELONGATED HYPOCOTYL 5 (HY5) and *B. distachyon* for bZIP16 but was excluded from the seed plant analysis. In seed plants, four photosynthesis related genes were identified as conserved. These transcription factors were the forkhead-type protein AT2G21530, GLK1, NUCLEAR FACTOR Y B3 (NFYB3), and PHYTOCHROME INTERACTING FACTOR 8 (PIF8). Eight transcription factors were enriched in all except one seed plant. Within these were several GCN5-RELATED N-ACETYLTRANSFERASEs (GNATs) including ATNSI, MYB SUPERFAMILY PROTEIN (MYBS1), B-BOX DOMAIN CONTAINING PROTEIN (BBX15), NRL PROTEIN FOR CHLOROPLAST MOVEMENT1 (NCH1), NUCLEAR FACTOR Y C4, (NF-YC4), and ZINC FINGER PROTEIN 1 (PNT1). Six transcription factors were eniched in all except two seed plants. Within these transcription factors were several GNATs and CONSTANS-LIKE 5 (COL5 also known as BBX6), GATA, NITRATE-INDUCIBLE, CARBON METABOLISM-INVOLVED (GNC), BELL1 (BEL1) and ARABIDOPSIS NAC DOMAIN CONTAINING PROTEIN 34 (ANAC034) (Figure 4A). These transcription factors belong to 12 families. GRN comparison between species can be used to identify conserved, high value targets for validation. Force-based drawing of photosynthesis target genes and their regulators using a Fruchtermann-Reingold algorithm (Fruchterman and Reingold, 1991) visualized the number of connections between TFs and between TFs and targets and is a different measure of centrality. By examining the network of *Z. mays*, we noticed that these conserved regulators (Figure 4A) were located in the centre of the network (Figure 4B) and this was also observed for the photosynthesis networks of seed plants (Figure S1). To quantify the observations, we examined the frequency of predicted regulated photosynthesis related genes. In the analysed species the conserved regulators regulate a higher percentage of photosynthesis related genes compared to the non-conserved transcription factors (Figure 4C).

**Figure 4:**
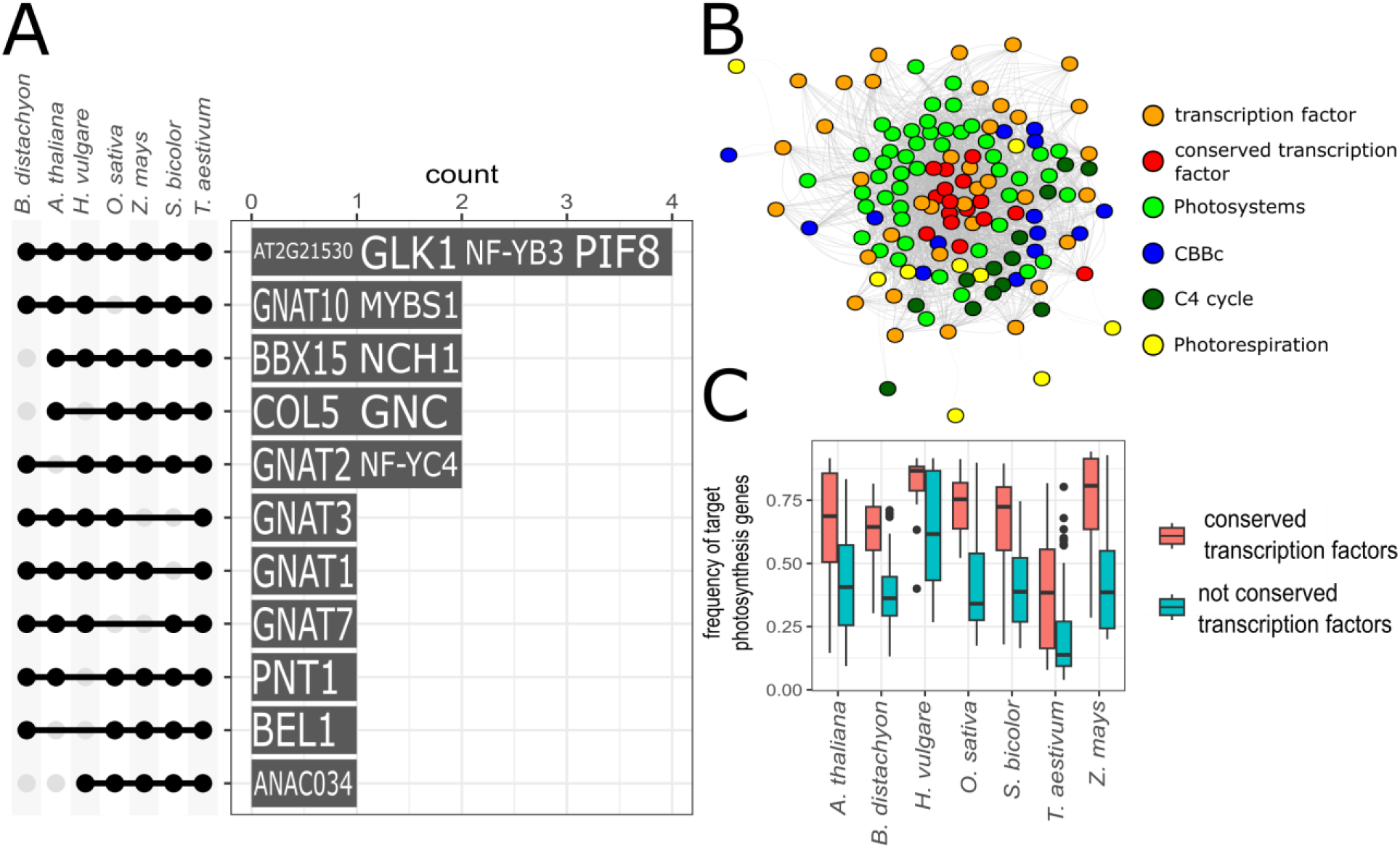
Identification of conserved photosynthesis regulators: A: Number of regulators with an enrichment in photosynthesis related genes in T. aestivum, S. bicolor, Z. mays, O. sativa, H. vulgare, A. thaliana, and B. distachyon with a common best Blast hit to A. thaliana with more than 4 species. B: Photosynthesis network of Z. mays. C: Frequency of predicted photosynthesis target genes of conserved transcription factors (>4 species).

Previous work showed that after WGDs, not all classes of genes are conserved alike (Almeida-Silva and Van de Peer, 2023). Therefore, we examined other agronomically relevant traits with the goal to identify potential targets for plant manipulation through breeding or genome editing. Especially in crop plants, generation of transgenic plants is a bottleneck. We wanted to demonstrate how the comparative network approach is able to narrow down candidates beyond sequence-based homology. We imagined the goal of increasing the identification of transcription factors involved in seed development in crops. To achieve that, a classical approach would be to investigate orthologue genes of a gene annotated for seed development in *A. thaliana*. Because of genome duplications, the number of orthologue genes in different species high. To reduce the number of possible candidates, we are now able to check if the predicted target genes of the transcription factors are conserved. This results in a better candidate selection process. Possible redundancy of transcription factors could be identified, which would require higher order mutants to observe a phenotype (Figure 5A). To demonstrate how the approach works, we selected the three *A. thaliana* transcription factors AT1G72570 (AP2-LIKE ETHYLENE-RESPONSIVE TRANSCRIPTION FACTOR, AIL1), AT1G77950 (AGAMOUS-LIKE 67, AGL67), and AT3G24650 (ABSCISIC ACID INSENSITIVE 3, ABI3) all of which have a GO term enrichment related to seed development. We then looked up the degree of conservation (Figure 2, Table S6, Figure 5B). AIL1 is involved in proper timing of seed filling in *A. thaliana* (Gao et al., 2011). AGL67 modulates seed germination under high temperature (Li et al., 2020). ABI3 is essential for seed maturation in *A. thaliana* (Delmas et al., 2013). For AIL1 only one orthologue with a high degree of conservation was identified for *S. bicolor* (p<1e-10). For the remaining grass species, orthologues were not conserved (Figure 5C). For AGL67, none of the orthologue transcription factors showed conserved target genes from *A. thaliana* to the grass species (Figure 5C). For ABI3, between *A. thaliana* and the grass species most orthologues were conserved in predicted function. *S. bicolor* had two orthologues for ABI3. One had a significant overlap in the target genes and the other did not. *T. aestivum* had three orthologues for ABI3, of which only two show a comparably high degree of conservation relative to the third orthologue. This similarity in conservation suggests that in *T. aestivum* a higher order mutant is necessary for a visible phenotype (Figure 5C). The approach is thus able to tease apart conserved from divergent evolution not only for pathways of primary metabolism (Figure 3) but also for regulation of seed development (Figure 5).

**Figure 5:**
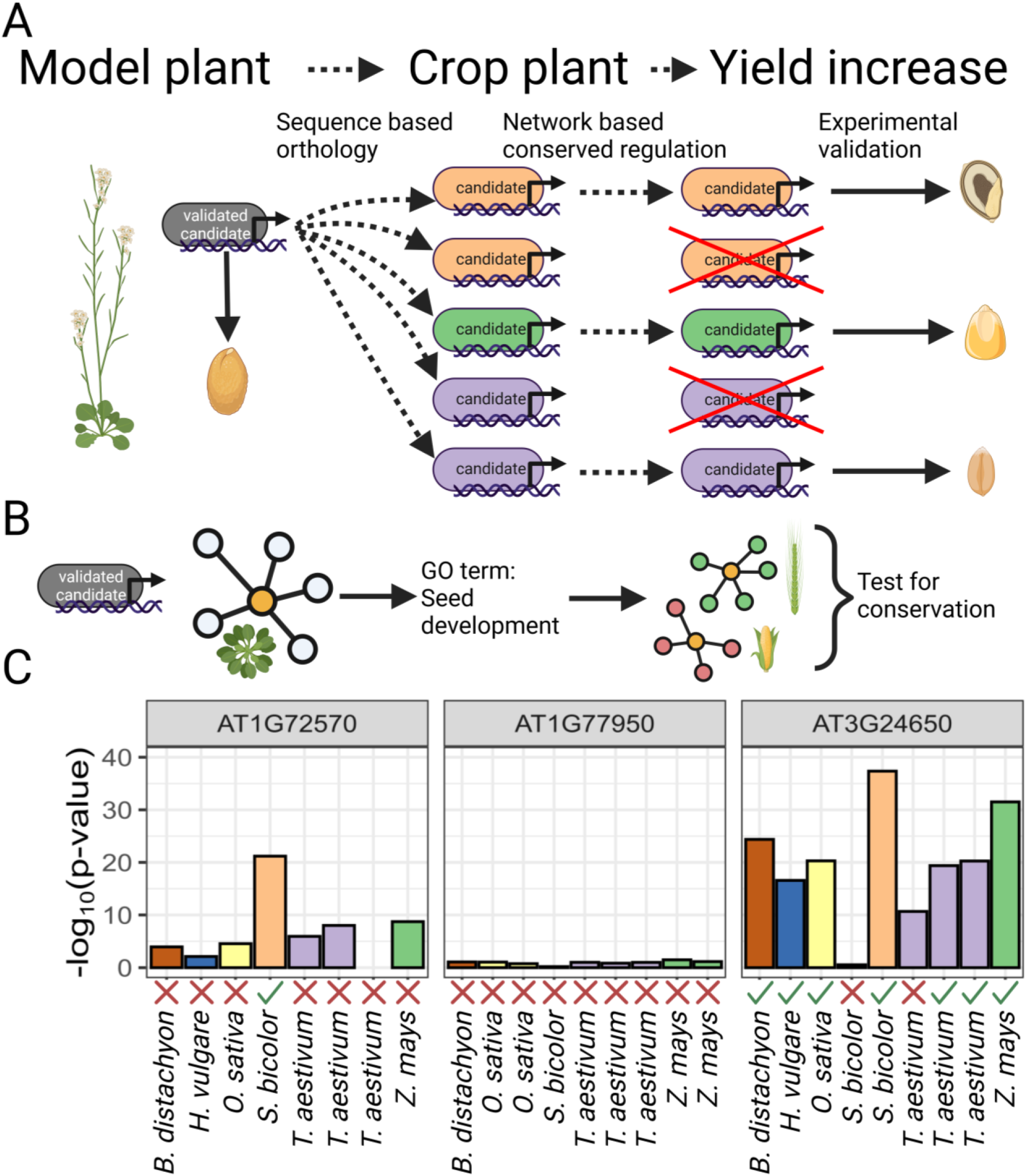
Identification of candidate genes across different species using combined orthologue and network approach. A: General concept how the approach can be used to identify transcription factors based on sequence and network based conservation. B: Specific example for seed development. C: Barplot of the log10 transformed p-values of the network based conservation of three A. thaliana candidates with an enrichment for seed development.

## Discussion

The gene regulatory network of *A. thaliana* and *C. reinhardii* were both validated by mutant and overexpressor analyses in previous studies (Halpape et al., 2023; Wulf et al., 2023). Therefore, they act as a good indicator if the network inference of the other networks was similarly successful. The frequency of TFs with a GO term enrichment was highest in *A. thaliana* but stayed above 33% for all species tested. This is in stark contrast to the different amounts of input data (Table 2) used for network calculations and indicates that the network approach is at least to some degree able to overcome differences in sampling and sample number (Table 1). The number of functions represented by the unique enriched and unique top enriched GO terms (Figure 1C and D) show that despite their differences, the GRNs contain similar amounts of biological information. It is likely that inadvertent variation introduced by different laboratories, e.g. sampling time, soil type, water and nutrient regimes, biotic contaminations in growth chamber, glass houses, and fields contribute to sufficient variation between data sets in the sequence read archives. The GRN with the highest sample number and medium genome complexity, *A. thaliana*, and the GRN with the second lowest sample number and very high genome complexity, wheat (Table 2), have similar information content (Figure 1 C, D). A likely reason for the high information content of the wheat GRN is the highly curated, specifically sampled, highly diverse dataset (Ramírez-González et al., 2018) used for its construction. Currently, no lower limit of sample number for successful construction can be established as the GRN information content also depends on the variation included in the dataset. The information content of the GRNs did vary from 30-50% of TFs with enrichment in a GO term (Figure 1A) and with 37 terms in *C. reinhardii* to 567 terms in *T. aestivum* processes represented (Figure 1D). This variation in information content indicates that the degrees of conservation estimated between networks are a lower bound of conservation. Increased sample number with increased diversity will likely increase information content, e. g in *B. distachyon*, as the other grasses with more samples or more variation included show higher information content (Table 2, Figure 1C, D). The GRNs are not without clearly observable limitations. The TF candidate identification for regulation of photosynthesis clearly showed the limitations of transcript abundance based GRNs. The known photosynthetic regulators PIF1, 3, 4, and 5 were not identified as conserved regulators in the GRN approaches (Figure 4A). These regulators are controlled to a large degree not by transcript abundance, but by post-translational modification, in this case phosphorylation (Shen et al., 2008), and by protein degradation via CONSTITUTIVE PHOTOMORPHOGENIC 1 (COP1) (Xu et al., 2017). As these processes are not captured in the transcript abundance based GRNs, these regulators are missed. For *C. reinhardtii*, the enrichment analyses with submodules of photosynthesis identified no candidate regulators for the CBBc and PR (Figure 3B). When mutants of TFs predicted to be relevant for photoprotection and for photosynthesis were analyzed, a role for a subset of TFs in regulating CBBc and PR was shown (Wulf et al., 2023). The use of a significance cut-off for candidate identification (p < 1e-10) may contribute to the problem. The examples show that for GRNs absence of evidence is not evidence for absence and therefore conclusions from absence need to be drawn with care, especially when cut-off based measures are used. Three different limitations of GRNs become evident. As described above, any TF regulated to a large degree on the posttranscriptional level will likely be missed by transcript abundance based GRNs. Processes which are not well described by a GO term are not captured. For plants, this includes, for example, the CBBc which is included in the “photosynthesis” GO term but is not described by its own GO term. Redox regulation is also not well captured in GO terms. And finally, conditions which are not represented in the dataset used for GRN construction, likely translate to poor annotation of the controlling TFs. The upper limit of half the TFs annotated with a likely function based on their targets in the GRNs may be close to the upper limit of possible annotations for the first two reasons.

With the knowledge of only being able to estimate a lower bound of conservation, the analyses proceeded. Three different viewpoints, from the dicot *A. thaliana*, from the monocot *Z. mays*, and from the alga *C. reinhardtii* were chosen for comparison. The dicot viewpoint revealed similar degrees of conservation with regard to observed p-values, frequencies and conserved processes towards all grasses reflecting the shared evolutionary distance of 150 million years between a dicot and monocots (Chaw et al., 2004). The much more distant alga with 950 million years to the last common ancestor (Magallón et al., 2013; Hedges et al., 2004) and a vastly different life style as a single celled organism (Boyes et al., 2001; Nishimura, 2010) has a much reduced degree of conservation. The viewpoint from an Andropogonae grass demonstrates highest conservation with the closest relative, *S. bicolor*, at least 50% significantly conserved TFs in all grasses with smaller values for the dicot and much smaller values for the alga (Figure 2). From the algal viewpoint, conservation is detectable but low for all seed plants (Figure 2). The conservation therefore perfectly recapitulates phylogeny. The processes for which conservation is detected down to the alga are those which represent general cellular functions such as protein translation (Table S6).

We used the well-defined process of photosynthesis to test if evolutionary changes in traits can be traced in the GRNs keeping the evidence caveat in mind. An eigenvalue-based analysis (Figure 3E) was added to avoid overconfidence in the threshold based measure (Figure 3A-D). During evolution towards the Andropogonae, the carbon concentration mechanism C4 photosynthesis was acquired (Sage, 2004). It has been hypothesized that this acquisition included the addition of the C4 cycle genes to the photosynthetic regulon (Külahoglu et al., 2014; Aubry et al., 2016). Indeed, both TF enrichment analyses which depend on thresholding via p-values (Figure 3C) and eigenvalue based centrality measures which use all available data (Figure 3E) show linkage which is not detectable in C3 species. The network suggests regulators of C4 photosynthesis are also regulators of the CBBc, and LHC&PS (Figure 3D). The presence of a complete C4 cycle reduces flux through the photorespiratory pathway (Sage, 2004) and reduces photorespiratory gene expression (Bräutigam et al., 2011, 2014; Bräutigam and Gowik, 2016). Indeed, the three species with CCMs show that photorespiration is at least partially unlinked from the photosynthesis module (Figure 3E). These analyses demonstrate that even with the limitations of GRNs based on sequence read data rather than dedicated data, evolutionary processes are reflected in the GRN comparison. It also demonstrates that the evolution or loss of a trait unlinks its regulation from its ancient position. The inferred regulatory networks differentiate between different species with different regulatory traits without relying on comparable expression datasets.

If TFs regulating a particular trait are sought, comparative GRN analyses provide several avenues to identify high probability candidate gene for further mutant and overexpressor based analyses. Of the 18 TFs identified as regulatory of photosynthesis in the comparative GRN analysis, several have been experimentally shown to control photosynthesis. Four TFs were predicted as photosynthetic in all GRNs: a fork-head domain protein (AT2g21530), GLK, NF-Y B3, and PIF8. For GLK-type TFs in Arabidopsis, tomato and rice mutants a downregulation of photosynthesis genes and a pale-green leaf phenotype was observed (Wang et al., 2013; Nguyen et al., 2014). Conserved transcription factor binding sites were identified near photosynthesis genes in *A. thaliana,* tomato, tobacco, maize and rice (Tu et al., 2022). Overexpression of NF-Y B3 alters photosynthetic gene expression in *T. aestivum* (TaNF-YB3, TraesCS3B01G017400) (Stephenson et al., 2011) and in *A. thaliana* (Halpape et al., 2023). In Arabidopsis, the PIF8 branch also contains PIF7 which is absent from grasses (Inoue et al., 2016) and PIF7 complementation lines in a *pif1,4,7* background restore photosynthetic gene expression (Halpape et al., 2023). PIF8 itself was shown to repress phyA-mediated light responses in *A. thaliana* (Oh et al., 2020). Of the six TFs identified in six out of seven GRNs, three are validated as photosynthetic. The myb-like transcription factor MYBS1 was initially described as a TF involved in sugar signalling (Chen et al., 2017). It regulates photosynthesis in *A. thaliana* (Halpape et al., 2023) and was found to be involved in the regulation of photosynthesis in the liverwort *Marchantia polymorpha* (Frangedakis et al., 2023). BBX15 acts downstream of GLK1 and GLK2 and both are direct targets (Susila et al., 2023). In *Petunia corollas* it was found that overexpression of BBX15 results in a higher chlorophyll accumulation (Ohmiya et al., 2019). NCH1 was shown to be involved in chloroplast accumulation in *M. polymorpha* and *A. thaliana* (Suetsugu et al., 2016). The NF-YC4 interacts with NF-YB3 and BBX proteins containing a CTT domain, namely BBX1 better known as CONSTANS (Lv et al., 2021). This opens the intriguing possibility of complex formation of NF-Y B3, NF-Y C4, and other BBX proteins of which NF-Y B3 and BBX14 (Halpape et al., 2023) and BBX15 (Lv et al., 2021; Susila et al., 2023) have been shown to control photosynthesis. Finally, the known photosynthetic regulator GNC in Arabidopsis (Chiang et al., 2012; Behringer and Schwechheimer, 2015) was identified in five out of seven GRNs. Its function is conserved in *Populus trichocarpa* (An et al., 2020). Several GNAT family transcriptional regulators were identified to be conserved to varying degrees. These GNATs are located in the chloroplast and some may alter photosynthetic activity by acetylation of photosynthesis proteins in *A. thaliana* (Ivanauskaite et al., 2023). For the remaining candidates, no information is known or other roles have been described. Similar to MYBS1, they may have dual roles and also control photosynthesis. COL5 is involved in flowering in *A. thaliana* (Hassidim et al., 2009). ZmBELL10 (Zm00001d035907) is the gene homologous to AtBEL1 and ZmBELL10 positively regulates growth and binds to genes involved in cell-division and elongation (Yu et al., 2023). ANAC034 regulates CONSTANS expression and has an effect on flowering time (Yoo et al., 2007). Photosynthesis may have a role in promoting flowering (Kandeler et al., 1975), so tests of these TFs may reveal a dual role or may show that at reduced levels of conservation, reliability of prediction is reduced.

Based on this we concluded that the network inference and evolutionary comparison is a useful tool for candidate identification for the regulation of biological processes. With our proposed workflow (Figure 5) we demonstrated how this comparative network approach is able to identify high confidence targets for mutant generation and validation. This validation step is a bottleneck for researchers, because of it being labour intensive and plant growth takes time (Kang et al., 2022). A network based prescreening of candidates can lead to faster allele identification and breeding for future yield increases.

## Author Contributions

D.W. designed and carried out computational analyses, interpreted data and co-wrote the paper.

A.B. conceived the initial idea and the study, interpreted data and co-wrote the paper.

## Acknowledgements

This work was supported by the BMBF-funded de.NBI Cloud within the German Network for Bioinformatics Infrastructure (de.NBI) (031A532B, 031A533A, 031A533B, 031A534A, 031A535A, 031A537A, 031A537B, 031A537C, 031A537D, 031A538A).

## Supplemental materials

All supplemental material are available on git (https://gitlab.ub.uni-bielefeld.de/donat.wulf/comp_net)

## Notes

https://gitlab.ub.uni-bielefeld.de/donat.wulf/comp_net

### Competing Interest Statement

The authors have declared no competing interest.

